# Predicting Functional Associations using Flanking Genes (FlaGs)

**DOI:** 10.1101/362095

**Authors:** Chayan Kumar Saha, Rodrigo Sanches Pires, Harald Brolin, Maxence Delannoy, Gemma Catherine Atkinson

## Abstract

Analysis of conservation of gene neighbourhoods over different evolutionary levels is important for understanding operon and gene cluster evolution, and predicting functional associations. Our tool FlaGs (<u>Fla</u>nking Gene<u>s</u>) takes a list of NCBI protein accessions as in input, clusters neighbourhood-encoded proteins into homologous groups using sensitive sequence searching, and outputs a graphical visualization of the gene neighbourhood and its conservation, along with a phylogenetic tree annotated with flanking gene conservation. FlaGs has demonstrated utility for molecular evolutionary analysis, having uncovered a new toxin-antitoxin system in prokaryotes and bacteriophages. FlaGs can be downloaded from https://github.com/GCA-VH-lab/FlaGs or run at www.webflags.se.

## Introduction

Conservation of gene order at long evolutionary distances is a strong indicator of a functional relationship among genes (Overbeek, et al. 1999). Extreme examples are the tryptophan biosynthesis (Dandekar, et al. 1998), and *str* ribosomal protein operons (Lechner, et al. 1989), which are conserved from bacteria to archaea. While such functional clustering is most commonly associated with prokaryotic genomes, it can even be observed in eukaryotic genomes (Lee and Sonnhammer 2003). The vast amount of genomic sequence data that has become available in recent decades is a treasure trove of clues about the function of uncharacterised proteins, and the pathways in which they are involved (Gabaldon and Huynen 2004). High-throughput identification of gene order conservation in genomes is a promising approach for predicting the involvement of proteins in particular pathways or systems. For example, gene neighbourhood conservation has been used in the prediction of novel proteins involved in adaptive immunity through association with CRISPR-Cas systems (Shmakov, et al. 2018). In addition to yielding functional predictions, the identification of conserved genomic architectures is essential for understanding the evolutionary dynamics behind the formation and restructuring of gene clusters, including reassembly of operons after disruption during evolution (Omelchenko, et al. 2003).

While there are a range of tools that analyse gene neighbourhood conservation or integrate this data along with other metrics for functional association prediction, these tend to be either restrictive in the genomes that can be considered (for example only complete genomes or those of model organisms) or require the creation of local genome databases (Lemoine, et al. 2008; Martinez-Guerrero, et al. 2008; Overmars, et al. 2013; Szklarczyk, et al. 2015; Garcia, et al. 2019). Other tools that connect to the NCBI (National Center for Biotechnology Information; https://www.ncbi.nlm.nih.gov/) to detect operons may lack sensitive sequence searching for homology assignments of neighbourhood genes (Gumerov and Zhulin 2020). We felt there was a need for a tool that allows use of the huge quantity of publicly accessible data in the NCBI RefSeq database (O’Leary, et al. 2016) and is sensitive enough to answer questions about homologous proteins over any evolutionary distance the user is interested in, from the strain or isolate level, to inter-kingdom or even inter-domain comparisons. We set out to build a Python tool that fulfils our list of essential criteria:

1. Allows the user complete control over the input genomes being analysed
2. Has a simple input format that does not require coding, downloading of genomes or formatting of databases
3. Nevertheless also has the option of running using locally stored genomes for offline analyses or analysing genomes that are not public
4. Can be run via a server with results emailed to the user
5. Can detect remote homology, suitable for analysing the most distant relationships between proteins and taxa as well as closer comparative analyses
6. Outputs gene neighbourhood annotated onto a phylogenetic tree
7. Produces publication-quality editable vector graphics.

## The FlaGs workflow

Our resulting tool that fulfils the above requirements is called FlaGs (standing for <u>Fla</u>nking <u>G</u>ene<u>s</u>) (**Fig. 1A**). FlaGs takes in user-determined NCBI accessions that link to the RefSeq database (around 170 million proteins from almost 100,000 organisms as of March 2020). Non-RefSeq NCBI protein accessions, or those that are out of date, are automatically updated to corresponding RefSeq accessions where possible. Input files can be easily and quickly prepared from selected sequences in the output of an NCBI BlastP search against the RefSeq database without any scripting (see the manual; Supplementary Materials file 1). An optional addition to the input file is the NCBI genome assembly identifier to target a particular genome. FlaGs clusters retrieved flanking gene-encoded proteins using the sensitive Hidden Markov Model-based method Jackhmmer, part of the HMMER distribution (Eddy 2011). There are three ways to run FlaGs:

1. Through the web server at www.webflags.se
2. Locally, with FlaGs querying NCBI as it runs, and not requiring locally stored genomes
3. Locally, using locally stored genomes in GFF format.

**Figure 1.**
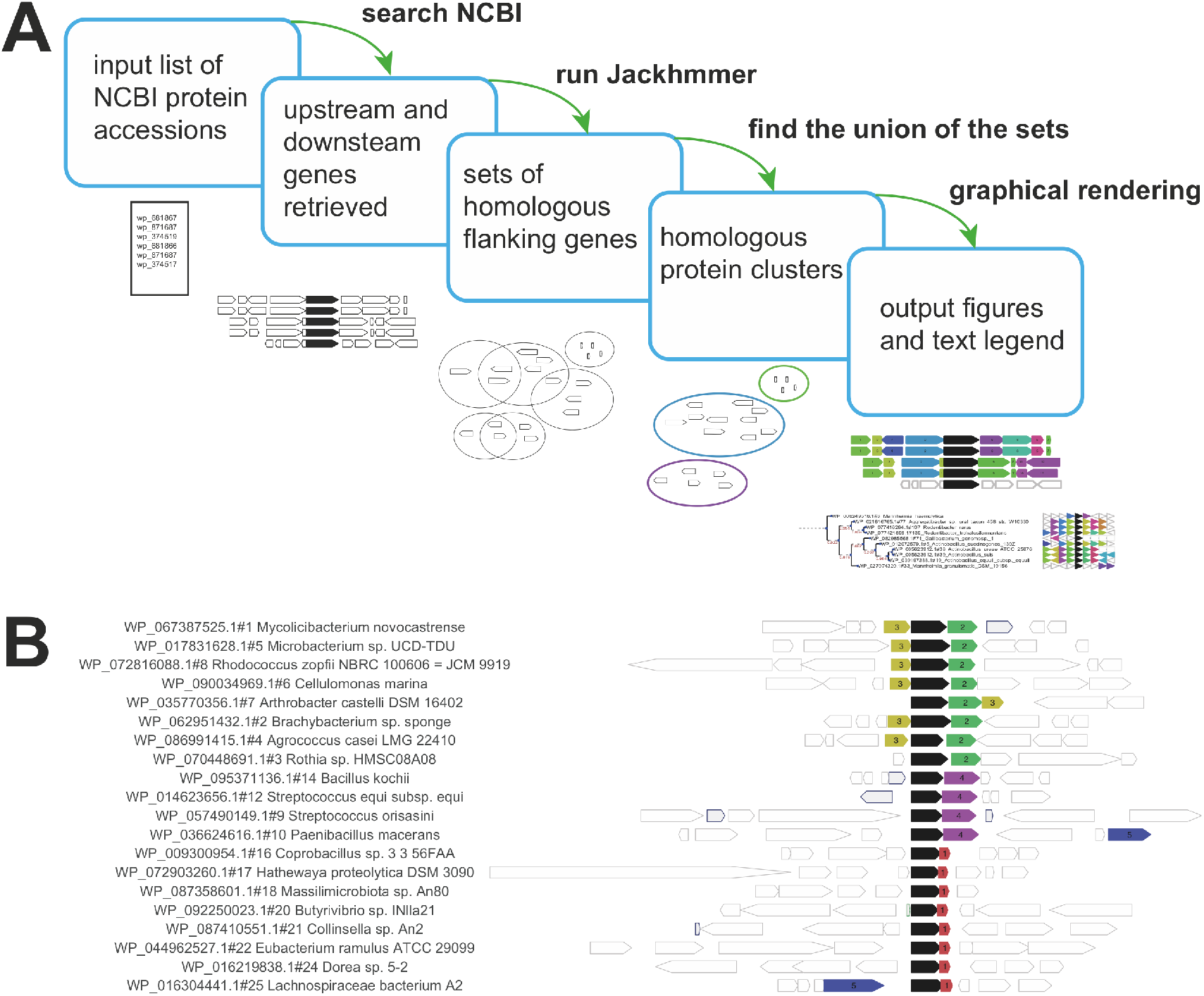
The FlaGs workflow and example results. (**A**) The user inputs a list of protein accession numbers – optionally with GCF assembly IDs – and can specify the number of adjacent flanking genes to consider, and the sensitivity of the Jackhmmer search though changing the E value cut-off and number of iterations. The output always includes a to-scale figure of flanking genes, a description of the flanking gene identities as a legend, and optionally, a phylogenetic tree annotated with colour- and number-coded pennant flags. (**B**) Example results using toxins of the toxSAS toxin-antitoxin system (Jimmy, et al. 2020) as the query. Empty genes with grey borders are not conserved in the data set, and grey genes with blue borders are pseudogenes. In this example, FlaGs reveals four different homologous groups of antitoxins as flanking genes, two of which (green and yellow) are antitoxins for the same cognate toxin. Group number 5 is an integrase. As FlaGs does not require complete genomes, regions can lack flanking genes on one side if the query gene is close to the end of a contig, as is the case with *Arthrobacter castelli* in this example.

FlaGs outputs information on the conservation of flanking gene-encoded proteins, and their identity, in graphical and text format (**Fig. 1A**). The output always includes a to-scale diagram of flanking genes, number- and colour-coded by conservation groups (**Fig. 1B**). A “description” file is also included, which acts as a legend for interpreting the flanking gene diagram. An optional output is a phylogenetic tree annotated with flanking genes reduced to triangular pennant-like flags. The tree-building feature uses the ETE 3 Python environment (Huerta-Cepas, et al. 2016).

FlaGs is a flexible tool for sensitive detection of flanking gene conservation at any evolutionary distance, and displays results in an intuitive, figure-quality graphical format. Possible applications of FlaGs include the discovery of novel functional associations and the analysis of gene neighbourhood dynamics during evolution. The utility of FlaGs is exemplified by our recent discovery of a novel toxin-antitoxin system exploiting growth control via ppApp alarmone nucleotide signalling (Jimmy, et al. 2020).

## Supporting information

Supplemental file: FlaGs manual

## Acknowledgements

We wish to thank Lars Barquist for suggesting the use of Jackhmmer in our pipeline, and Marek Wilczynski for help setting up the WebFlaGs server. The work was supported by Vetenskapsrådet (the Swedish Research Council; grants 2015-04746 and 2019-01085 to GCA), and UCMR Linnaeus Program Gender Policy Support (to GCA).

