## Supplemental file: FlaGs manual for "Predicting Functional Associations using Flanking Genes (FlaGs)"

### FlaGs User Manual
### May 2020

#### https://github.com/GCA-VH-lab/FlaGs/
#### Server: webflags.se

**Chayan Kumar Saha**


**Gemma C. Atkinson**


**Contents:**

**1. Description** page 2
**2. Prerequisites and use of the easy install script** page 4
**3. Usage** page 6
**4. Running the example files** page 6
**5. Arguments** page 7
**6. Output files** page 10
**7. Running online with WebFlaGs** page 12
**8. Recommendations and tips** page 13
**9. References** page 14

**Appendix** page 14

Creating an input file for FlaGs from an NCBI BlastP or PSI-Blast search

### 1. Description

FlaGs (for Flanking Genes) analyses genomic context around genes encoding a protein of interest using a Python3 script combined with Jackhmmer (Eddy 2011) and, optionally, ETE-toolkit (Huerta-Cepas, et al. 2016). FlaGs finds the flanking genes upstream and downstream of the genes of interest, clusters them based on homology and represents them visually with distinct identifiers (colour and number).

At initiation the tool verifies the input file which is either a list of NCBI protein accessions (-p option), or a tab delimited list of both protein accessions and corresponding genome assembly identifers (-a). The latter option is to inform FlaGs which specific genome it should be searching with the protein accession; since NCBI protein accessions are non-redundant, identical protein sequences can be present in multiple genomes with the same accession number. If FlaGs is given protein accessions alone using the –p option, it takes each accession and finds the corresponding list of assembly identifiers (eg. *GCF_000001635.23 or GCA_000001635.5)*. Then it can either report all results for all identifiers (-r option), or for one representative identifier (default). The –r option should be used with caution as identical proteins can sometimes be found in thousands of genomes from closely related species, strains or isolates. See *Arguments* below for guidance.

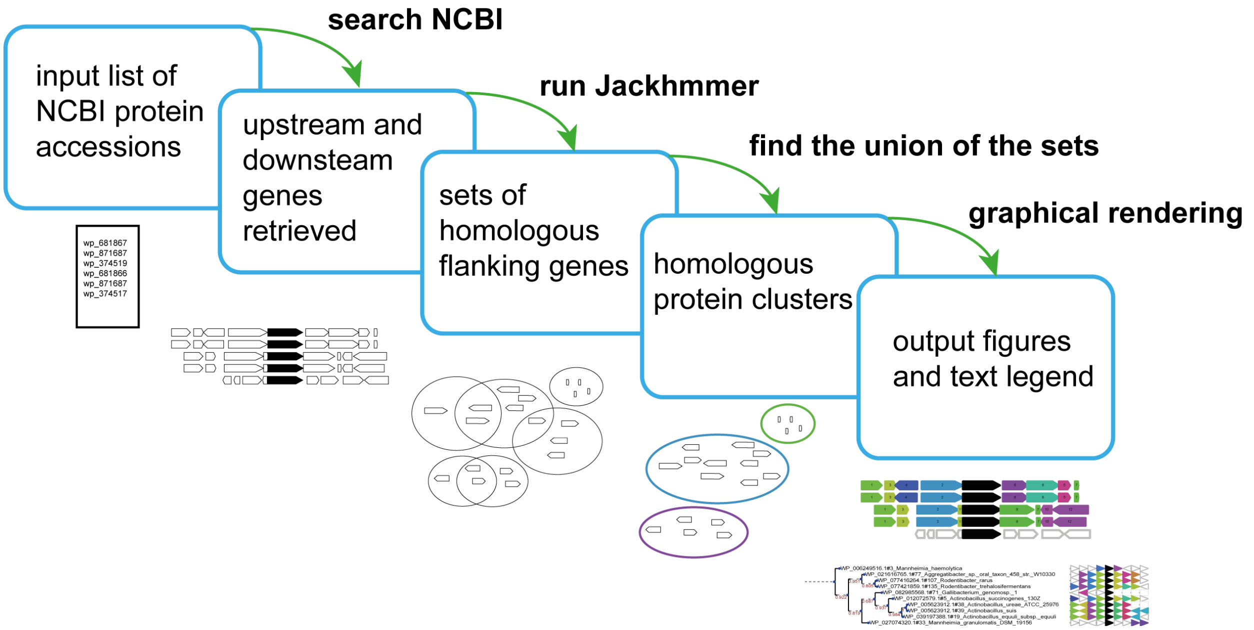


**Figure 1. The FlaGs workflow.** The user inputs a list of protein accession numbers – optionally with GCF assembly identifiers – and can specify the number of adjacent flanking genes to consider and the sensitivity of the Jackhmmer search though changing the E value cut-off and number of iterations. The output always includes a to-scale figure of flanking genes, a description of the flanking gene identities as a legend, and optionally, a phylogenetic tree annotated with colour- and number- coded pennant flags. Unconserved proteins are uncoloured.

After verification of the input query list, FlaGs uses the assembly identifier to retrieve the Genomic Feature Format (gff3) file for each protein query and uses this to identify annotated flanking genes from upstream and downstream regions. Then it retrieves the sequences for all flanking genes and searches for homologues within the set using Jackhmmer from the Hmmer package (hmmer.org, (Eddy 2011)). Flanking genes are then clustered and each cluster is assigned a specific number. Sets of homologues found with the Jackhmmer searches are joined together to make one cluster (the union) if there is one or more common protein among the Jackhmmer search results (Figure

1). The numbering of clusters begins with 1. The lower the cluster number the more conserved the protein is, i.e the more frequently it is found among all the flanking genes. Finally, FlaGs generates a visual representation of the output using the tkinter Python module, and this is saved as a postscript file (Figure 2). The black block arrow represents the query proteins and the rest represent the flanking genes. The figure is proportional to actual gene lengths and intergenic space. If the query protein is encoded on the negative strand, the entire context is then flipped to make it easily comparable. The number and colour represent the cluster, and the "…_outdesc.txt" output file provides the legend for interpretation. Unconserved proteins are uncoloured and unnumbered: , RNA coding genes are outlined in green: , and pseudogenes are grey, outlined in dark blue:

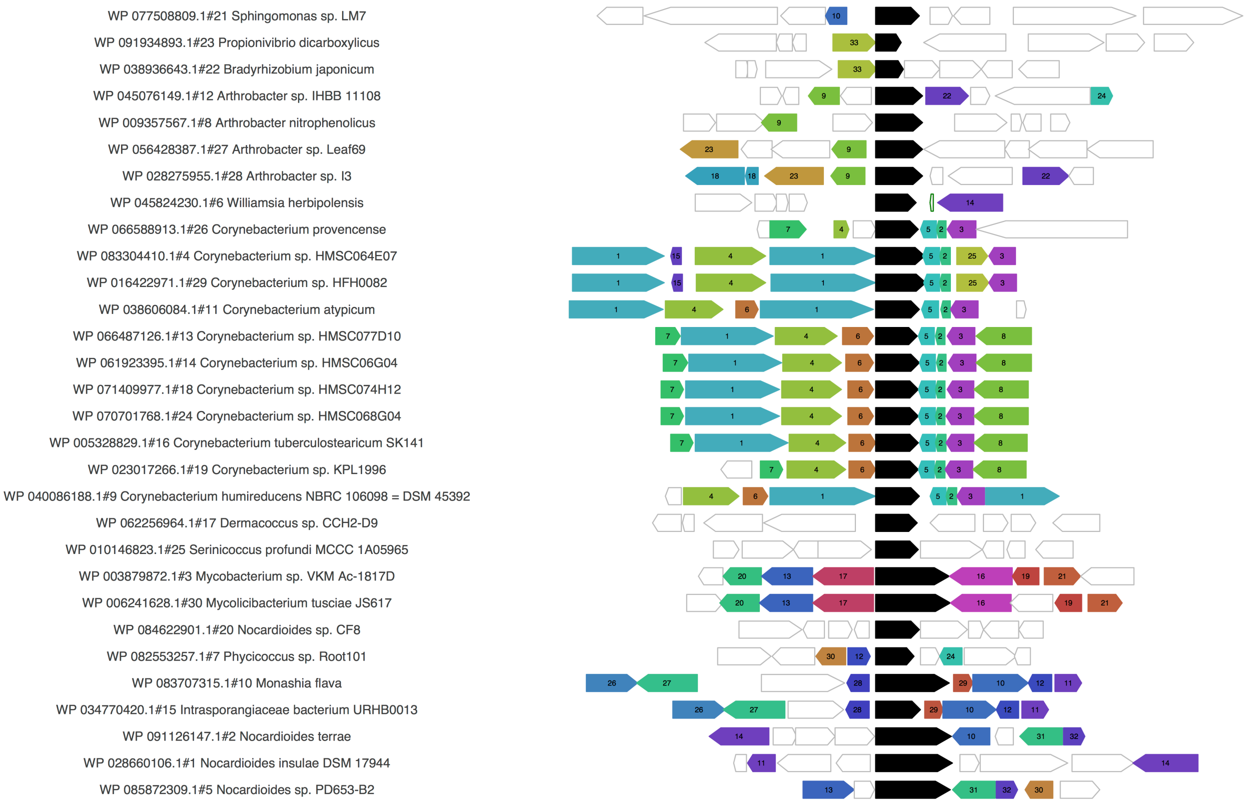


**Figure 2. The to-scale postscript output.** The black gene is the query, and otherwise colours and numbers represent the clusters. The "…_outdesc.txt" output file provides the legend for interpretation. Genes and intergenic spaces are to scale.

An optional feature of FlaGs is the generation of a phylogenetic tree of the query sequences, on which the flanking genes are annotated (Figure 3). This is achieved using the ETE-toolkit (etetoolkit.org, (Huerta-Cepas, et al. 2016)). The ETE "mafft_default-trimal01-none-fasttree_full" workflow is used to generate the tree (see ETE user manual, http://etetoolkit.org/cookbook/). The flanking genes and the queries are represented as triangular pennant flag-like shapes, with colours and numbers representing clusters. The "…_outdesc.txt" output file again provides the legend for interpretation.

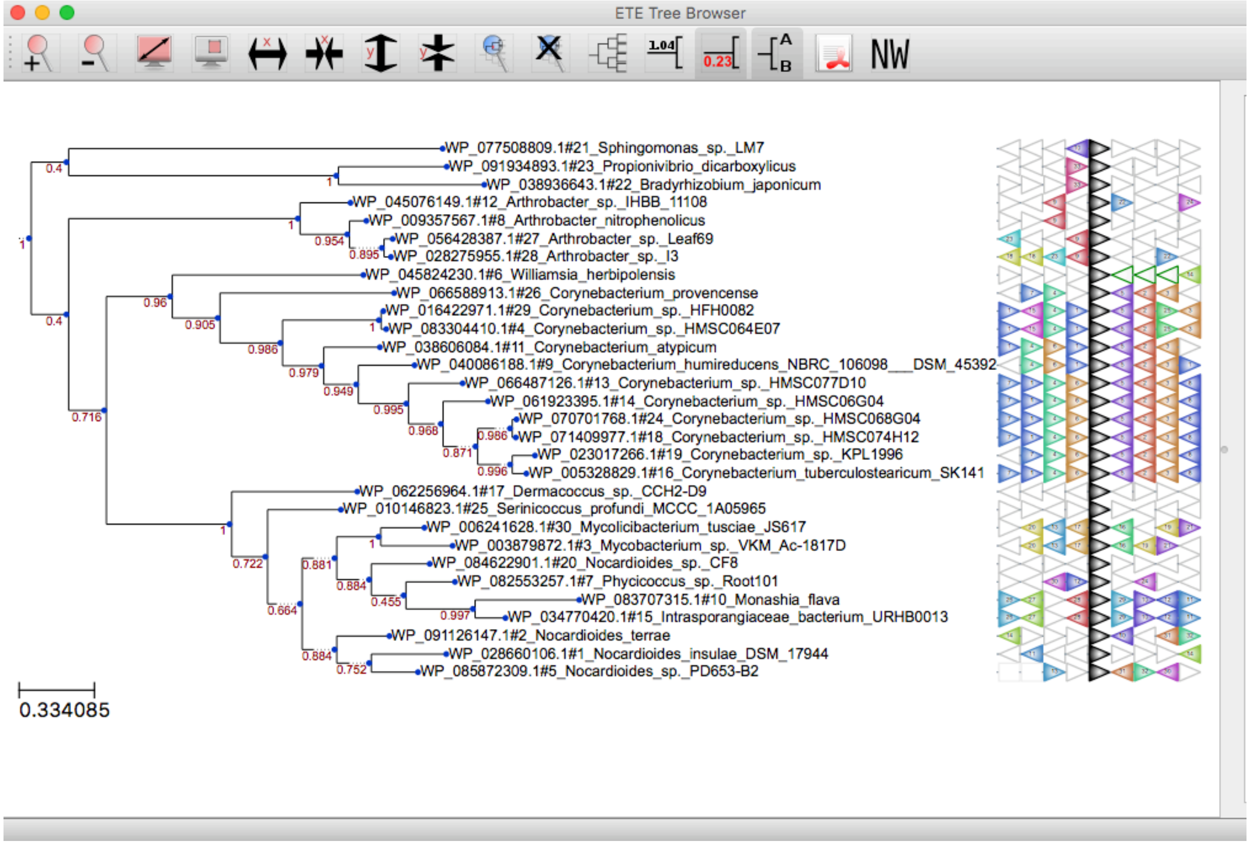


**Figure 3. The annotated phylogenetic tree output.** The tree is made with ETE3 using the mafft_default-trimal01-none-fasttree_full method and is saved as an SVG vector image file. If the –t option is invoked without –to, an interactive ETE3 window opens (as in this figure) for manipulation of how the tree is displayed.

#### 2. Prerequisites and use of the easy install script

FlaGs.py is available for download here: [https://github.com/GCA-VH-lab/FlaGs/](https://github.com/GCA-VH-lab/FlaGs/)

FlaGs has been primarily tested in Mac and Linux environments. Users have reported that running on Windows is possible with some workarounds (updates in this direction are planned). In the meantime, Windows users are directed to use the server version at webflags.se instead.

For Mac and Linux users, a bash script **install.sh** is bundled with the download, which installs the programs required for running FlaGs. If you are using a Mac and do not yet have Xcode command line tools installed, you will be prompted to do so, following the on-screen instructions. Navigate to the FlaGs folder and use the following command to run the quick install script (answering y or yes when required):

```
bash ./install.sh
```

If this does not work due to insufficient permissions, and you have an admin/root password you can use sudo (But! Always safety first when using sudo: by all means check the install.sh file first to make sure you're comfortable with the commands that will be run (the list of tools that will be installed follow below)):

```
sudo bash ./install.sh
```

Tools that will be installed by the script (if they are not already installed):
Homebrew (to install Wget)
Wget (to install Anaconda)
Anaconda (to install the tools below)
Hmmer
Biopython
ETE (which itself installs a number of phylogenetic tools)

Alternatively, users can install the prerequisite programs manually following the instructions below:

1. Python 3

We recommend the use of Anaconda Python to simplify installation of other required and optional dependencies. It can be downloaded from https://www.anaconda.com/download/ If this works and you don't want to install a specific Anaconda version, you can proceed to step 2.

Recommendation for specific Anaconda version:

Anaconda3 (version 5.2.0) is well tested with FlaGs. It is available for both Linux and Mac Users in the anaconda archive (Link https://repo.anaconda.com/archive/). The following command can be used to download anaconda 5.2.0.

#For Mac users (wget required)

```
wget https://repo.anaconda.com/archive/Anaconda3-5.2.0-MacOSX-x86_64.sh
```

#For Linux users

```
wget https://repo.anaconda.com/archive/Anaconda3-5.2.0-Linux-x86_64.sh
```

2. Biopython

Using Anaconda, Biopython can be installed using the following command:

```
conda install -c conda-forge/label/gcc7 biopython
```

3. Jackhmmer

Using Anaconda, hmmer can be installed using the following command:

```
conda install -c biocore hmmer
```

Hmmer can alternatively be downloaded from http://hmmer.org/ and installed according to the instructions.

4. ETE-toolkit (only required if making a phylogenetic tree; –t option)

### Install ETE and external tools
```
conda install -c etetoolkit ete3 ete_toolchain
```

### Check installation
```
ete3 build check
```

### Activate the environment (you can add the line to your .bashrc to make it permanent)
```
export PATH=~/anaconda_ete/bin:$PATH
```

(these instructions come from http://etetoolkit.org/download/)

### 3. Usage:

Mac OS:
```
./FlaGs.py  <options>
```
or `Python3 FlaGs.py <options>`
or (if Python 3 is the default python) `Python FlaGs.py <options>`

Linux:
replace `FlaGs.py` above, with `FlaGs_linux.py`

For options and arguments, see below.

### 4. Running the example files:

N.B. a valid email address is required by NCBI to monitor the number of requests to their server per second per user. You will not receive emails from the local version of FlaGs.

Without tree (only Biopython neccessary):

```
python3 FlaGs.py -a GCF_accession_input.txt -o GCF_accession_output -u -vb
```

or

```
python3 FlaGs.py -p accession_input.txt -o accession_output -u -vb
```

With tree (Biopython and ETE necessary):

```
python3 FlaGs.py -a GCF_accession_input.txt -t -to -o GCF_accession_output -u -vb
```

or

```
python3 FlaGs.py -p accession_input.txt -t -to -o accession_output
-u -vb
```

With tree and interactive ETE tree editing window:

```
python3 FlaGs.py -a GCF_accession_input.txt -t -o
GCF_accession_output -u -vb
```

or

```
python3 FlaGs.py -p accession_input.txt -t -o accession_output -u
 -vb
```

### 5. Arguments:

1. "-a", "--assemblyList"

Protein accession with Genome Assembly Identifier eg. GCF_000001765.3 in a text input file separated by a tab.

Input File Example:

GCF_000001765.3    WP_047256880.1 #tab separated
GCF_000002753.1    WP_012725678.1
…….

2. "-p", "--proteinList"

Protein Accession WP_047256880.1 in a text Input file separated by newline. If these are not RefSeq accessions (WP_), the program will attempt to convert to a RefSeq accession.

Input File Example:

WP_047256880.1
WP_012725678.1
…….

3. “-l”, “--localGenomeList”

FlaGs v1.0.5 can use local genomes too. For that, the user needs to provide the following files:

i. Compressed (gzip) FASTA format of the predicted protein products annotated on the genome assembly (eg. Assembly_1.faa.gz, file should contain “.faa.gz” suffix)

An example of the recommended format can be downloaded from this link. Or use the following command from the terminal:

```
wget
ftp://ftp.ncbi.nlm.nih.gov/genomes/all/GCF/000/428/725/GCF_
000428725.1_ASM42872v1/*ftp://ftp.ncbi.nlm.nih.gov/genomes/
all/GCF/000/428/725/GCF_000428725.1_ASM42872v1/GCF_00042872
5.1_ASM42872v1_protein.faa.gz
```

ii. Compressed (gzip) Annotation of the genomic sequence(s) in Generic Feature Format Version 3 (GFF3) (eg. Assembly_1.gff.gz, file should contain the ".gff.gz" suffix)

The recommended format can be downloaded from this link. Or use the following command from the terminal:

```
wget
ftp://ftp.ncbi.nlm.nih.gov/genomes/all/GCF/000/428/725/GCF_
000428725.1_ASM42872v1/GCF_000428725.1_ASM42872v1_genomic.g
ff.gz
```

With the "-l" flag set, the user specifies the input text file with assembly ID and the query protein accession.

Input file example:

GCF_000001765.3    WP_047256880.1 #tab separated
GCF_000002753.1    WP_012725678.1

In this case, both of the Fasta and gff file should contain the Assembly Identifier (the GCF number) as the prefix.
For example, for assembly identifier GCF_000001765.3, the compressed Fasta file should be named GCF_000001765.3.faa.gz and the GFF file should be named GCF_000001765.3.gff.gz

Or input file example with user-defined assembly names:

Assembly_1    WP_047256880.1 #tab separated
Assembly_2    WP_012725678.1

For assembly identifier Assembly_1, the compressed Fasta file should be named as Assembly_1.faa.gz and the GFF file should be named as Assembly_1.gff.gz

4. "-ld", "--localGenomeDirectory"

This flag allows the user to specify the path or directory where *.faa.gz* and *.gff.gz* are stored. By default, FlaGs script will look for the *.faa.gz* and *.gff.gz* files in the same directory where the script is located or running from.

The user can only use this flag when the '-l' flag is being used.

For example: If the files are stored in ~/FlaGs/local directory user can specify by following:

```
-ld ~/FlaGs/local
```

5. "-r", "--redundant"

This will initiate a search for flanking genes in all available GCFs that encode this identical protein sequence (using -r A or -r a) for each query, or a specific number of GCFs (for example to search a maximum of 5 genomes encoding identical protein sequences, the user would write -r 5).

Warning: the –r A option should be used with caution as identical proteins can sometimes be found in thousands of genomes in RefSeq. Using -r with a number to limit is highly recommended.

6. "-e", "--ethreshold"

This e-value is used by Jackhmmer as a cutoff parameter to detect homology among all flanking genes. By default it is 1e-10.

7. "-n", "--number"

This number represents number of Jackhmmer iterations that allows to find more distant homolog, by default it is set as 3.

8. "-g", "--gene"

Using this parameter, the user can define the number of upstream and downstream genes to look for each query in the input list. By default, it is 4, which means for each query it will try to find 4 upstream and 4 downstream flanking genes and then process further.

9. "-t", "--tree"

This requires an ETE3 installation. The option enables showing flanking genes along with a phylogenetic tree.

10. "-ts", "--tshape"

This requires an ETE3 installation and thus this option only works when -t is used. This parameter can increase or decrease the size of triangle shapes that represent flanking genes, by default it is 12.

11. "-tf", "--tfontsize"

This also requires an ETE3 installation and thus this option only works when -t is used. This parameter can increase or decrease the size of font inside triangles that represent flanking genes, by default it is 4.

12. "-to", "--tree_order"

In combination with –t, it will first generate the tree output, and then use the tree order to generate the other (without tree) output file.

This is very useful option for stitching the two output files together in (for example) Illustrator to make a figure with both the tree, and the to-scale neighbourhood output.

13. “-u”, “--user_email”

A valid email address is required by NCBI to monitor the number of requests to their server per second per user. (there is a limit of 3 queries per second, which may be exceeded if you run multiple instances of FlaGs in parallel; but see the API-key advice below). You will not receive emails from the local version of FlaGs.

14. “-api”, “--api_key”

Valid API-key allows 10 queries per seconds, which makes the tool run faster. This may sometimes allow the user to run multiple instances of FlaGs in parallel, but in general this is not well tested or advised.

For details on the API key, please check:

https://ncbiinsights.ncbi.nlm.nih.gov/2017/11/02/new-api-keys-for-the-e-utilities/

15. "-o", "--out_prefix"

Any Keyword to define your output eg. MyQuery. This will appear in the name of your output files.

16. "-k", "--keep"

If the user wants to keep the intermediate files eg. gff3, this option is used (warning: these are large files). By default, intermediate files will be removed.

17. "-v", "--version"

Retrieve the version number of the program.

18. "-vb", "--verbose"

This option will show the work progress for each query as STDOUT. This is a very useful option, which we recommend using.

### 6. Output files

In addition to the figure output files described in *1. Description* above, FlaGs generates the following output files:

1. Information about the flanking genes for each of the queries retrieved from databases is stored in a tab delimited file named with "_flankgene.tsv" suffix
2. A general summary report is also generated for each query, for example if the query accession is valid or invalid, if the query accession was converted or updated to another accession associated with an identical RefSeq sequence, if the query failed, or if flanking gene information is found or not. This is a tab delimited file with the "_QueryStatus.txt" suffix
3. A general flanking gene summary report is also generated for each valid query if flanking gene information is found or not in a tab delimited file with "_flankgene_Report.log" suffix
4. A file with the suffix "_jackhits.tsv" contains the *Jackhmmer* output
5. After the clustering steps a file named with the suffix "_clusters.tsv" is generated with detailed information of the cluster. For example;

   1 4 WP_001229260.1;WP_001229255.1
   2 4 WP_001164213.1;WP_001164217.1;WP_001164219.1;WP_055027268.1

   Column 1 represents the cluster number; the lower the number the more frequent it is in the figure. The second column shows the frequency of proteins in the cluster. In first row (Cluster 1), 4 means that two protein accessions WP_001229260.1 and WP_001229255.1 are 4 times frequent and in cluster 2 you can see 4 proteins accessions are frequent for 4 times.

6. A description file is generated after clustering, with the suffix "_outdesc.txt". **This is the main legend file for interpreting the figure files.** For example;

   1(2) WP_001229260.1 MULTISPECIES: integration host factor subunit alpha
   1(2) WP_001229255.1 MULTISPECIES: integration host factor subunit alpha

   2(1) WP_001164213.1 MULTISPECIES: phenylalanine--tRNA ligase subunit alpha
   2(1) WP_001164217.1 MULTISPECIES: phenylalanine--tRNA ligase subunit alpha
   2(1) WP_001164219.1 phenylalanine--tRNA ligase subunit alpha
   2(1) WP_055027268.1 phenylalanine--tRNA ligase subunit alpha

   Here you can see in column 1, it starts with cluster number 1 (this is the number shown within genes in the figure files) and "(2)" means the protein WP_001229260.1 has been found twice in the cluster and the last column is the protein description retrieved from the database. The protein description gives information about the proteins in a cluster (accessions and title from NCBI).

   If wanted, this file can be used as an input for the additional bundled python script descriptionCloud.py to save a wordcloud visualization. descriptionCloud.py has additional dependencies: pandas, PIL, wordcloud and matplotlib. If these are not already installed, they are available through anaconda, eg:
   ```
   conda install -c conda-forge wordcloud
   ```

   descriptionCloud.py is run as follows:

   ```
   python descriptionCloud.py -i example_outdesc.txt
   ```

7. Discarded protein ids with improper accession are listed in a text file with "_NameError.txt" suffix.
8. Discarded protein ids lacking proper information in RefSeq DB are listed in a text file with "_Insufficient_Info_In_DB.txt" suffix
9. If the –t option used to make a phylogenetic tree, ETE3 saves additional results files to a folder ending with "[output name]_tree_". This includes a Newick format version of the tree.

FlaGs also generates some intermediate files to proceed the overall process but only the files mentioned above are useful for interpretation. Intermediate files can be kept with the -k option (warning: these are large files).

### 7. Running online with WebFlaGs

FlaGs can be run through the web server at www.webflags.se. WebFlaGs is a more user friendly version of FlaGs (as it does not require installation) with a web interface where user can paste the input in the specific text area or upload as text file. WebFlaGs accepts input datasets that contains up to 500 accessions. If the user wants to run more than 500 queries, it is recommended to use the local version of FlaGs. FlaGs and webFlaGs both use the same input file format.

In webFlaGs, the user can set the e-value cut-off that is used by Jackhmmer to detect homology among all flanking genes. By default, it is 1e-10. The user can also choose the number of Jackhmmer iterations that allows it to find more distant homologs. By default this is set as 3. Like FlaGs, in webFlaGs the user can define the number of upstream and downstream genes to look for flanking each query in the input list. By default, it is 4.

When running webFlaGs, a valid email address is required, which will be used for sending results. The user will receive a download link for a zipped file which contains:

1. The one or two figure files described above based on the output type selected in the webFlaGs webpage by the user. There are three output type options in webFlaGs. If the option 'With phylogenetic tree and query showed as tree order' is selected, the user will get the annotated phylogenetic tree output and the to-scale postscript output where queries will be presented as the same order that of the annotated phylogenetic tree output. For option 'With phylogenetic tree but query showed as input order', queries in the to-scale postscript output will be presented as the same order as submitted by the user, also the user will get the annotated phylogenetic tree output in svg format. If 'No phylogenetic tree and query showed as input order' is selected, the user will get only the to-scale postscript output along with the converted PDF file (see below).

2. A converted PDF version of the to-scale postscript output. This is provided for Windows users who can not easily open postscript files.

3. A description file, which is generated after clustering with suffix "_outdesc.txt" (described above). This is the main legend file for the figures.

4. The query input file with the suffix "_query.flagsIn"

5. A text file with the suffix "_QueryStatus.txt" suffix that records the success (or not) of each query, and importantly informs if the input accession failed but was converted or updated to an identical RefSeq sequence that worked.

6. (Optional)

   If the user selected 'With phylogenetic tree' as Output type option, only then the will get the multi-fasta sequence file with suffix "_tree.fasta" which was used for making tree file and the resulting Newick format version of the tree, having a suffix ".nw".

WebFlaGs uses the Entrez Programming Utilities (E-utilities) API to access the NCBI database system and retrieve the Genomic Feature Format (gff3) file for each protein query. After May 1, 2018, NCBI has limited this access, which is why WebFlaGs can't place more than 10 requests/second to NCBI. As a result, this usually makes the results processing a bit slower than with the local version of FlaGs.

### 8. Recommendations and tips

When the results are received, it is recommended that users check the tab delimited results file with the "_QueryStatus.txt" suffix. This records the success (or not) of each query, and importantly inform if the input accession failed but was converted or updated to an identical RefSeq sequence that worked. When using the -p option, the query status file is also useful for identifying which genome (identified with a GCF number) was used to retrieve the flanking genes.

When using the -to option, the two output figures show input taxa listed in the same order (by the tree). Therefore, in Adobe Illustrator (or similar) it is very easy to collate these two vector images together, in order to make a figure of the to-scale flanking gene output mapped on to the tree.

FlaGs can handle relatively large input datasets, although of course the smaller, the faster. Since FlaGs retrieves information from NCBI (multiple times for each input accession), it is dependent on both a stable local internet connection and the stability of the NCBI server. Our example input file "GCF_accession_input.txt" with 30 accessions takes around 7 minutes to run with the –a, -t, -to options on a 2015 MacBook Pro with an internet connection of around ~75 mbps. Without the -t option it takes around 6 minutes. We have also tested input files containing around 1000 protein accessions (four flanking genes each side) and they worked successfully over the course of a few hours.

If you have >1000 entries, you may want to reduce the number of input accessions. One way to do this is by reducing redundancy with a tool such as Usearch (Edgar 2010).

The -vb (verbose) option is very useful for following the progress of the run, and we recommend its use unless you want to limit what is displayed via STDOUT.

FlaGs is designed for use with prokaryotic and bacteriophage genomes where gene clusters can be conserved over vast evolutionary distances, and the space between genes is relatively small. While it will run with eukaryotic nuclear genomes, the running time is much longer and the resulting images can be very wide. For this kind of data, we recommend initial testing with small datasets and few flanking genes (the -g option). FlaGs does however work very well with eukaryotic organellar genomes that are more bacteria-like in their organisation and conservation.

If wanted, the _outdesc.txt file can be used as an input for the additional bundled python script descriptionCloud.py to save a wordcloud visualisation. descriptionCloud.py has additional dependencies: pandas, PIL, wordcloud and matplotlib. If these are not already installed, they are available through anaconda, eg:

```
conda install -c conda-forge wordcloud
```

descriptionCloud.py is run as follows:

python descriptionCloud.py -i example_outdesc.txt

A png format output image will be created, for example:

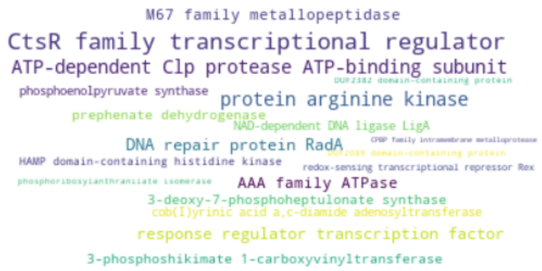

#### Appendix: Creating an input file for FlaGs from an NCBI BlastP or PSI-Blast search

##### 1. Run the search against RefSeq proteins:

Go to https://blast.ncbi.nlm.nih.gov/Blast.cgi?PAGE=Proteins , paste in your query sequence and select the refseq_protein database. Set any advanced parameters and/or organism limits you like, then click Blast.

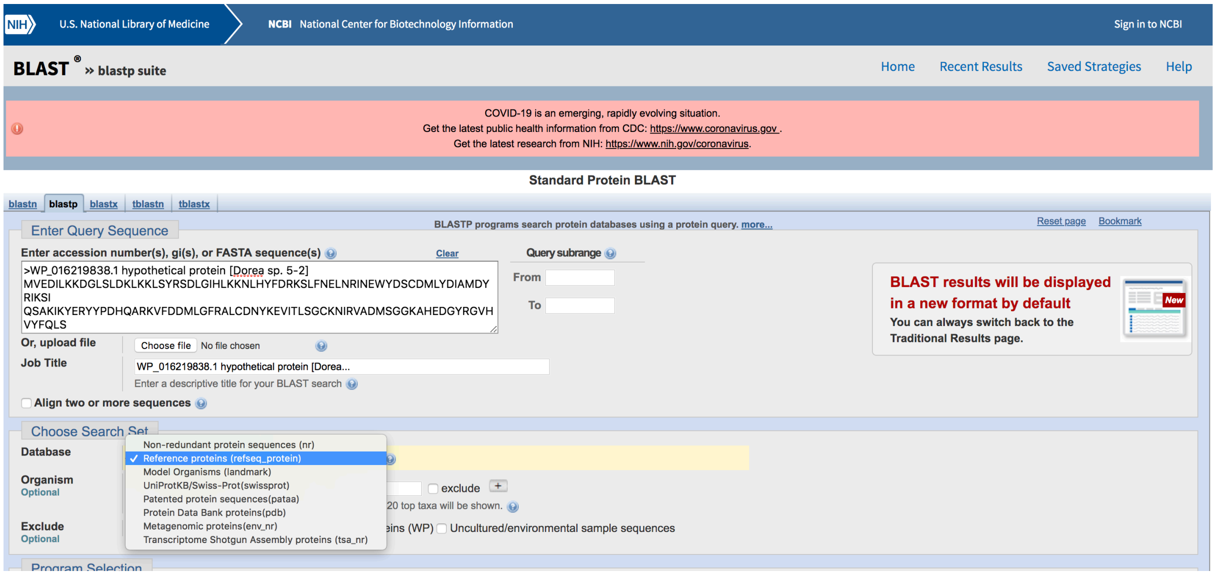


##### 2. When the results appear, select the proteins of interest

Use the check boxes next to each hit to select the results you're interested in, or leave "select all" checked.

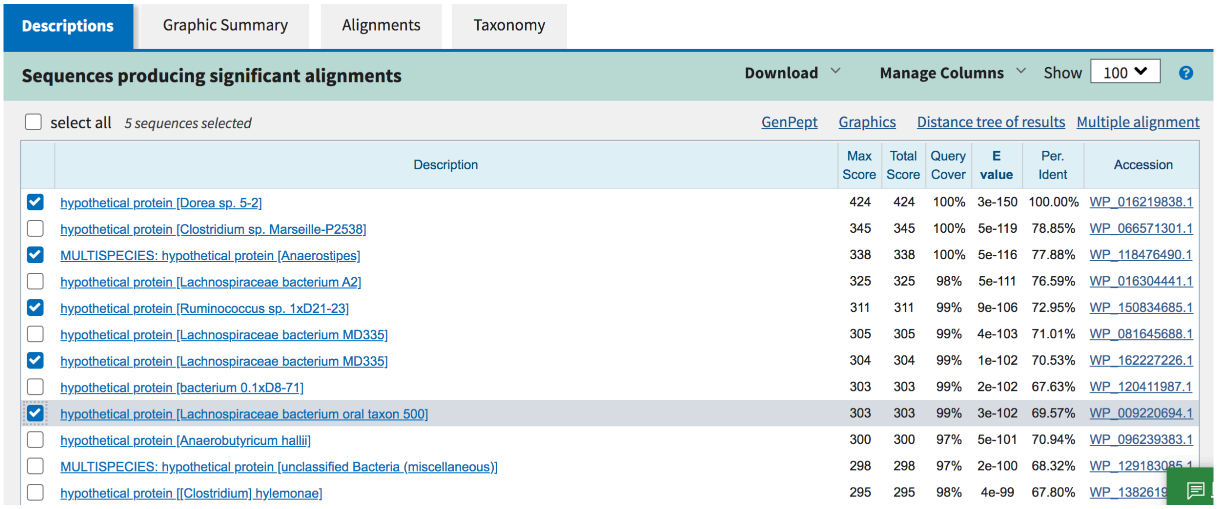


| | Description | Max Score | Total Score | Query Cover | E value | Per. Ident | Accession |
|---|---|---|---|---|---|---|---|
| ☑ | hypothetical protein [Dorea sp. 5-2] | 424 | 424 | 100% | 3e-150 | 100.00% | WP_016219838.1 |
| ☐ | hypothetical protein [Clostridium sp. Marseille-P2538] | 345 | 345 | 100% | 5e-119 | 78.85% | WP_066571301.1 |
| ☑ | MULTISPECIES: hypothetical protein [Anaerostipes] | 338 | 338 | 100% | 5e-116 | 77.88% | WP_118476490.1 |
| ☐ | hypothetical protein [Lachnospiraceae bacterium A2] | 325 | 325 | 98% | 5e-111 | 76.59% | WP_016304441.1 |
| ☑ | hypothetical protein [Ruminococcus sp. 1xD21-23] | 311 | 311 | 99% | 9e-106 | 72.95% | WP_150834685.1 |
| ☐ | hypothetical protein [Lachnospiraceae bacterium MD335] | 305 | 305 | 99% | 4e-103 | 71.01% | WP_081645688.1 |
| ☑ | hypothetical protein [Lachnospiraceae bacterium MD335] | 304 | 304 | 99% | 1e-102 | 70.53% | WP_162227226.1 |
| ☐ | hypothetical protein [bacterium 0.1xD8-71] | 303 | 303 | 99% | 2e-102 | 67.63% | WP_120411987.1 |
| ☑ | hypothetical protein [Lachnospiraceae bacterium oral taxon 500] | 303 | 303 | 99% | 3e-102 | 69.57% | WP_009220694.1 |
| ☐ | hypothetical protein [Anaerobutyricum hallii] | 300 | 300 | 97% | 5e-101 | 70.94% | WP_096239383.1 |
| ☐ | MULTISPECIES: hypothetical protein [unclassified Bacteria (miscellaneous)] | 298 | 298 | 97% | 2e-100 | 68.32% | WP_129183085.1 |
| ☐ | hypothetical protein [[Clostridium] hylemonae] | 295 | 295 | 98% | 4e-99 | 67.80% | WP_1382619[illegible] |

##### 3a: If you have <100 sequences selected, you can click "Genpept"

This takes you to a multi-protein summary page. You can change the format to "Accession list" (making sure the number of results shown per page is set high enough).

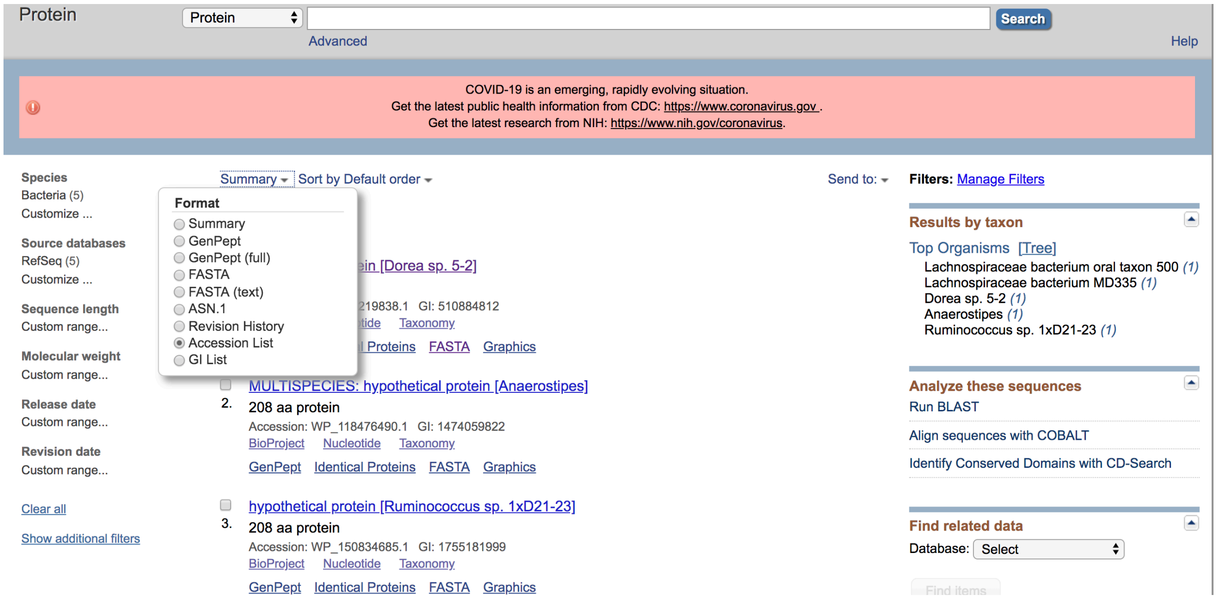


The list of accessions can be pasted into a text file to use as FlaGs input. You're done and ready to run FlaGs! You can stop reading here.

```
WP_016219838.1
WP_118476490.1
WP_150834685.1
WP_162227226.1
WP_009220694.1
```

##### 3b: If you have >100 sequences selected (or just want to try a different way), you can click "Download", then "Hit Table (CSV)"

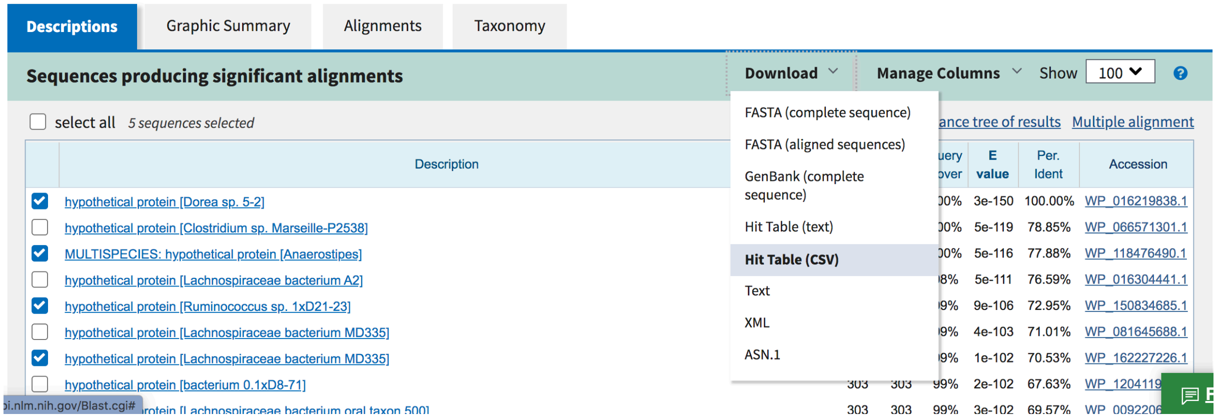


This is a table that can be opened in a spreadsheet program such as Excel, Numbers or Google Sheets

##### 4. Open your table, and split the text into columns

In Excel and Google Sheets this is done through the Data > Text to columns menu option

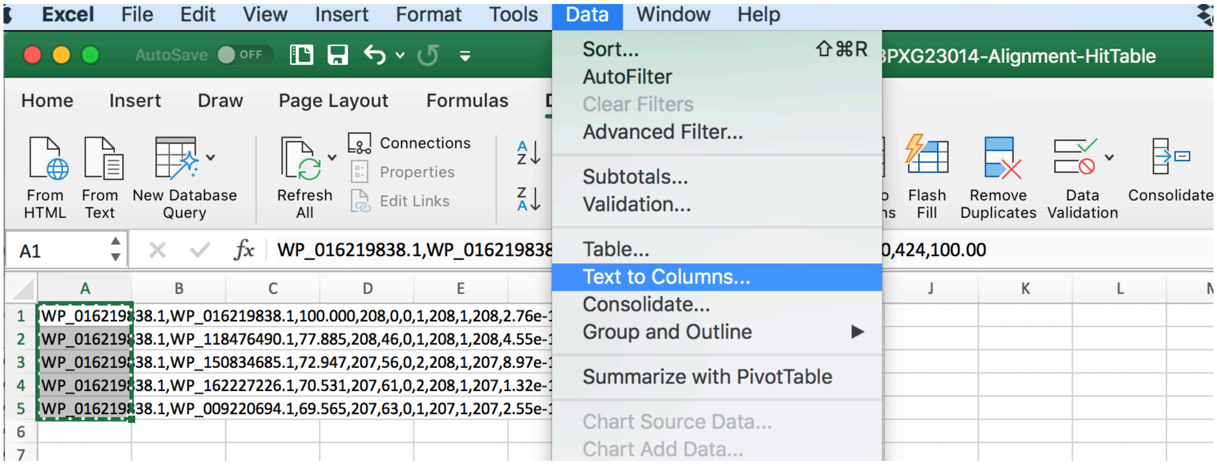


Follow the wizard to split by commas

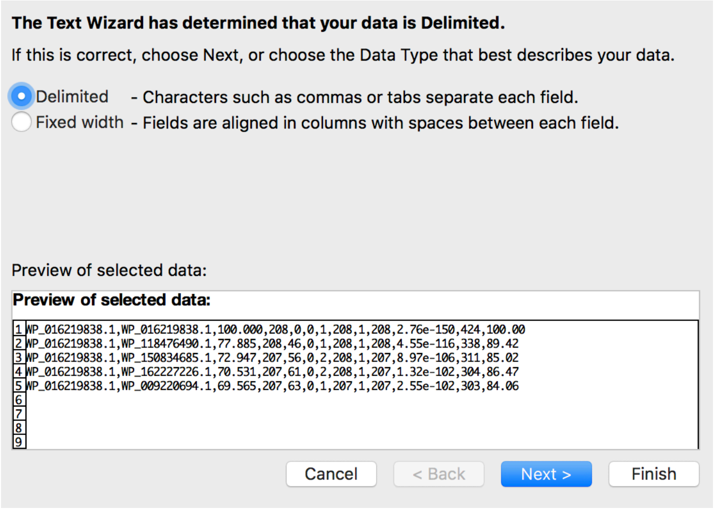


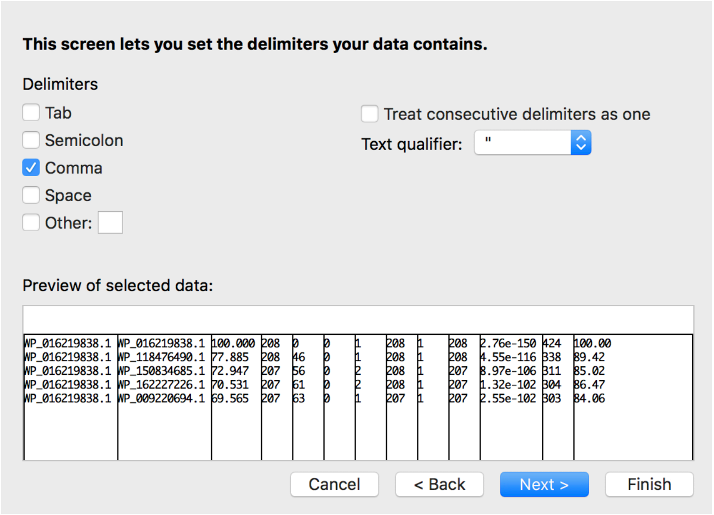


Click finish and you will see your list of accessions in the second column

| | A | B | C | D | E | F | G | H | I | J | K | L | M |
|---|---|---|---|---|---|---|---|---|---|---|---|---|---|
| 1 | WP_016219 | WP_016219 | 100.000 | 208 | 0 | 0 | 1 | 208 | 1 | 208 | 2.76e-150 | 424 | 100.00 |
| 2 | WP_016219 | WP_118476 | 77.885 | 208 | 46 | 0 | 1 | 208 | 1 | 208 | 4.55e-116 | 338 | 89.42 |
| 3 | WP_016219 | WP_150834 | 72.947 | 207 | 56 | 0 | 2 | 208 | 1 | 207 | 8.97e-106 | 311 | 85.02 |
| 4 | WP_016219 | WP_162227 | 70.531 | 207 | 61 | 0 | 2 | 208 | 1 | 207 | 1.32e-102 | 304 | 86.47 |
| 5 | WP_016219 | WP_009220 | 69.565 | 207 | 63 | 0 | 1 | 207 | 1 | 207 | 2.55e-102 | 303 | 84.06 |
| 6 | | | | | | | | | | | | | |

The list of accessions can be copied and pasted into a text file to use as FlaGs input. You’re done and ready to run FlaGs!

```
WP_016219838.1
WP_118476490.1
WP_150834685.1
WP_162227226.1
WP_009220694.1
```
